# Fractals in Action: An fMRI study on the Generation of new Hierarchical Levels in Motor Sequences

**DOI:** 10.1101/232942

**Authors:** Mauricio J.D. Martins, Roberta Bianco, Daniela Sammler, Arno Villringer

## Abstract

Generation of hierarchical structures, such as the embedding of subordinate elements into larger structures, is a core feature of human cognition. *Discrimination* of well-formed hierarchies is thought to rely on lateral prefrontal cortex (PFC). However, the brain bases underlying the *active generation* of new hierarchical levels remain poorly understood. Here, we created a new motor paradigm to isolate this *active* generative process. In fMRI, participants planned and performed (identical) movement sequences based on three previously learned rules: (1) a hierarchical ‘fractal’ rule that involved generation of new levels, (2) a linear ‘iterative’ rule adding items to existing hierarchical levels, and (3) simple ‘repetition’. We found that generation of new hierarchical levels (using the fractal rule) activated a bilateral motor planning-and imagery network, but did not involve lateral PFC. Conversely, adding items to existing hierarchical levels required M1 directly during execution. These results show that the generation of new hierarchical levels can be achieved without involvement of putative domain-general systems such as those ascribed to lateral PFC. We hypothesize that these systems might be important to *parse* hierarchical sequences in a multi-domain fashion but not necessarily to *generate* new hierarchical levels.

## Introduction

Much of what differentiates human behaviour from that of other species is related to an increased ability to represent and generate complex hierarchies (Fitch and Friederici 2012; Conway and Christiansen 2001; Everaert et al. 2015; Dehaene et al. 2015). Language, music and complex action planning are examples of human behaviours which require the representation and generation of hierarchies. The assumed centrality of this capacity to mankind has made the investigation of the underlying neural mechanisms a core topic in modern cognitive sciences (Badre 2008; Fadiga, Craighero, and D’Ausilio 2009; Fitch and Martins 2014; Friederici et al. 2011; Petersson, Folia, and Hagoort 2012; Wilson, Marslen-Wilson, and Petkov 2017). Numerous studies in the domains of language, music, and vision have investigated the *discrimination* of hierarchical structures by asking participants to evaluate whether sequences of items are well-formed according to a previously learned system of rules (Bahlmann et al. 2009; Bahlmann, Schubotz, and Friederici 2008; Bianco et al. 2016; Maess et al. 2001; Sammler, Koelsch, and Friederici 2011; Friederici et al. 2011). From this body of literature it emerges that lateral prefrontal cortex (PFC, particularly Inferior Frontal Gyrus, IFG) might contribute domain-general resources to the processing of hierarchies (Patel 2003; Friederici et al. 2011; Fadiga, Craighero, and D’Ausilio 2009), in interaction with areas along ventral visual/auditory streams that may store domain-specific structural and schematic information (Oechslin, Gschwind, and James 2017; Martins et al. 2014; Pallier, Devauchelle, and Dehaene 2011; Sammler et al. 2013; Bianco et al. 2016).

Despite the appeal and success of this framework, it remains incomplete in one essential way: It is currently unknown to which extent we can use these data from *discrimination* paradigms to understand the processes underlying the *generation* of hierarchical structures. This subtle distinction between *discrimination* and *generation* becomes more relevant when borrowing concepts from modern theories of hierarchical generativity in language (Berwick and Chomsky 2015), which propose that representing hierarchies involves (i) core generative processes, (ii) peripheral sensorimotor systems (for encoding and externalization), and (iii) interfaces between (i) and (ii). This framework implies that when the stimulus sets to be discriminated vary not only in their deep (hierarchical) structure but also in their surface (linear) structure (e.g. long-distance vs. local dependencies (AABB vs. ABAB)), they might impose different encoding and externalization demands due to peripheral parameters such as stimulus frequency, ambiguity, working memory load, etc.

In the present study, we investigated the *generation* of hierarchically organised behaviours and their neurocognitive bases by means of a novel paradigm in the motor domain where the generation of structures can be directly and explicitly measured at the behavioural level. Crucially, to circumvent differences related to the peripheral processes of stimulus encoding, our design experimentally separated the generative act from the externalization of structures which were derived from different rules but were identical in their surface (linear) structure, i.e. the sequential motor output.

So far, behavioural and neural markers of action production have been extensively studied typically in contexts of (i) fast production of movement sequences without hierarchical relations (Hardwick et al. 2013; Elsinger, Harrington, and Rao 2006; Hétu et al. 2013) and (ii) representation of hierarchical relations within action sequences but without generation of new hierarchical levels (Fazio et al. 2009; Koechlin and Jubault 2006).

The first stream of research is based on classical serial reaction time tasks (SRTT), based on the execution of sequences of items characterised by simple adjacent dependencies. This research has demonstrated that the learning of motor sequences is supported by a network including premotor and motor cortices (PMC and M1), superior parietal lobe (SPL) supplementary motor area (SMA), pre-SMA, left thalamus and right cerebellum (Hardwick et al. 2013). Furthermore, “planning” (defined as keeping a given motor sequence in short-term memory between the time of cue presentation and the time of execution) recruited a bilateral network comprising the sensorimotor and premotor cortices, cerebellum and the basal ganglia (Boecker et al. 2008; Elsinger, Harrington, and Rao 2006).

The second stream of research has focused on the neural bases supporting the hierarchical processing of actions (when certain acts are encoded as part of higher-order actions) (Fazio et al. 2009; Koechlin and Jubault 2006). These studies typically compare higher vs. lower levels of given and fixed hierarchical action structures and have revealed posterior-to-anterior activation gradients along lateral PFC with increasing hierarchy level. For instance, sequences of simple finger movements (left and right button presses) activate more anterior regions in lateral PFC when the movements are organized in superordinate clusters compared to un-clustered movement sequences (Koechlin and Jubault 2006). Some authors (Badre and D’Esposito 2009; Badre 2008) extrapolated these findings to everyday actions and hypothesize that when certain actions (e.g. slicing bread) occur within higher-order contexts (e.g. making a sandwich) these are encoded more anteriorly in lateral PFC in comparison to when the movement occurs without such higher-order action goal.

Overall, these studies provide neural evidence for (i) generative capacity but restricted to linear motor sequences and (ii) hierarchical processing of actions, but without generation of new hierarchical levels. The present study combined these two approaches to go a step further and probe the mechanisms underlying the internally driven generation of hierarchically organised motor sequences. Delineating the operational limits of different generative procedures (linear and hierarchical) is essential to uncover the neural bases specifically supporting the generation of motor hierarchies. More broadly, it may become critical to our understanding of how we represent generative rules in other domains, such as language and music.

Our approach involved the comparison of brain activity during active generation of motor sequences that follow different rules (‘Iterative’, ‘Fractal’) which can be used to generate or transform hierarchies (Martins 2012; Hulst 2010; Martins, Martins, and Fitch 2015). The first rule is called ‘Iterative’ and adds items linearly within levels of a given hierarchy without generating new levels. For example, with the rule “add 1 B subordinate to A”, we can start from [A[B]] and generate A[BB]], [A[BBB]], and so forth. The second rule is called ‘Fractal’ and allows the generation of new hierarchical levels. For instance, with the rule “add 2 α subordinate to α” we start with [α] and generate the hierarchies [α[αα]], [α[α[αα] α[αα]]], and so forth. These fractal principles can be described with a recursive notation [α] **→** [α [αα]], as in Lindenmayer systems (Lindenmayer 1968). This kind of re-write rules have been used to describe supra-regular grammars in linguistic syntax (Fitch and Friederici 2012), as well as visual (Martins, Martins, and Fitch 2015), melodic (Martins et al. 2017) and rhythmic fractals (Geambasu, Ravignani, and Levelt 2016), while experimental evidence for the use of these rules during explicit production of new hierarchical levels and their neural instantiation is still lacking.

In this fMRI study, we isolate how participants generate new hierarchical levels by contrasting Iterative and Fractal rules. Importantly, we do so in production (not discrimination) of well-formed hierarchies. Moreover, we use a procedure that separates the core generative act from its encoding/externalization components: In our task, participants performed sequences of finger movements evolving in three consecutive steps according to (1) the linear Iterative rule, (2) the hierarchical Fractal rule, or (3) simple Repetition of the preceding step. While participants executed the movements of the first two steps (I and II) that established the rule following visual cues, they freely generated the last step (III) by applying the rule themselves, without visual support. By focusing on the transition between steps II and III, we were able to isolate the neural representations involved in the generation of new hierarchical levels. Simple Repetition allowed us to extract a non-generative baseline.

From the literature reviewed above, two hypotheses emerge: (1) the generation of new hierarchical levels requires the recruitment of systems thought to be involved in the processing of well-formed hierarchies in action and across domains (with special focus on lateral PFC and IFG); or (2) this capacity can be supported by domain-specific networks known to instantiate motor operations. Our results speak in favour of the second hypothesis.

## Methods

### Participants

20 healthy participants (11 males and 9 females, age range 21-35, *M* = 26.5) took part in the study. All participants were non-musicians. None had more than 2 years of music training, and none practiced regularly with a musical instrument. All had normal or corrected-to-normal vision and audition, no history of neurological or psychiatric disease. All participants were right-handed German native speakers. Participants were recruited from a pool of subjects able to perform all behavioral tasks successfully (see below). They gave written informed consent before the experiment in accordance with the local ethics committee and were paid 8 Euros/hour for their participation.

### Task and Stimuli

Participants were asked to produce sequences of finger movements on a keyboard with 16 keys. They had to press a correct set of keys in the correct order, and with the correct timing. The keyboard contained visual and tactile markers on keys 3, 5, 7, 10, 12 and 14 (from left-to-right) for spatial reference (see Figure 1). The temporal structure (see Figure 1A) was given by an auditory metronome sounding at 60bpm (4 beats per second delivered through MR-compatible headphones).

**Figure 1.**
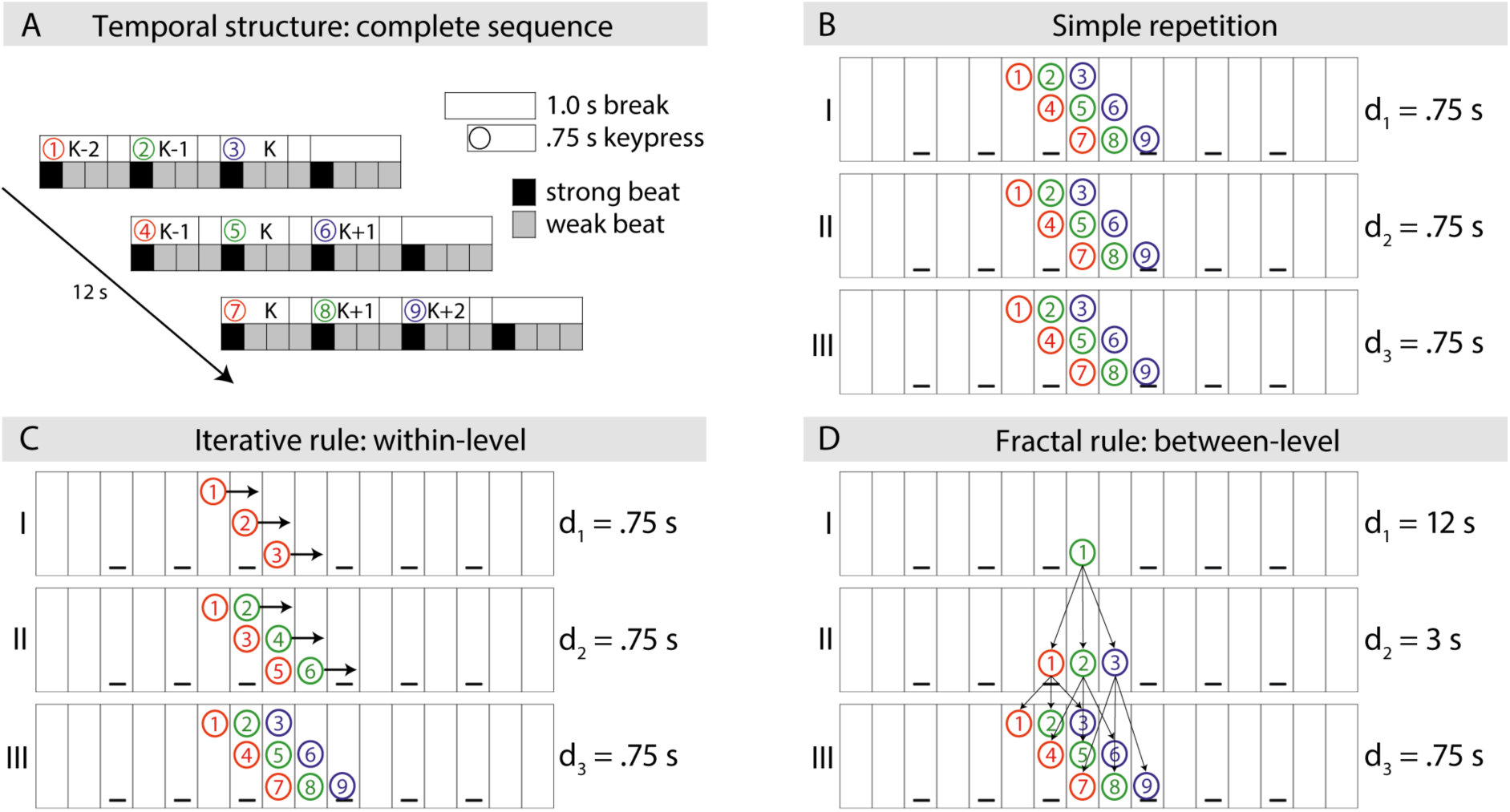
Task principles. In this task, participants were asked to generate sequences of finger movements (ordered from 1 to 9) by pressing keys on the keyboard with the thumb, index and middle finger (red, green and blue). These sequences were formed in three steps (I,II and III) which followed one of three rules: Simple Repetition, Iteration and Fractal. During steps I and II, participants executed the motor sequence as displayed on the screen. In step III, they were asked to generate the final sequence without visual support. ***A)Temporal structure:*** *In step III, all rules resulted in the same complete sequence of 9 movements, here [[K-2, K-1, K], [K-1, K, K+1], [K, K+1, K+2]]. K is the key in the spatial center of the pattern. The sequence lasted 12 seconds and was aligned with a metronome with 4 beats per second (1 strong and 3 weak). Key presses started at the onset of the strong beat, and were released at the onset of the third weak beat, thus, each key press had the duration (d) = .75s. Hierarchical clustering within the sequence (3 clusters of 3 items) was given by the fingering pattern (red, green and blue) and the temporal structure (1s break after each cluster).* ***B) Simple Repetition:*** *Consisted of the repetition of the complete sequence 3 times.* ***C) Iterative rule:****Step I was composed of 3 key presses executed with the thumb, each with d = .75s, on the first (strong) beat of each cluster [[K-2, _, _ ], [K-1, _, _ ], [K, _, _ ]. In step II, a second key press with the index was added to each chunk: [[K-2, K-1, _ ], [K-1, K, _ ], [K, K+1, _ ]]. Thus, the iterative rule added elements to pre-existing hierarchical levels, without generating new levels. Step III was simply the serial completion of the pattern with the middle finger [[K-2, K-1, K], [K-1, K, K+1], [K, K+1, K+2]].* ***D) Fractal rule:****Step I was a single key press with the index finger on key K with d = 12s. Step II was a sequence of three key presses [K-1, K, K+1] executed with the thumb, index, and middle finger, each with d = 3s and 1s break after each key press. The underlying fractal rule was the substitution of each key press K_n_ with duration d_n_ with a sequence of three key presses [K_n+1_ -1, K_n+1_, K_n+1_ + 1], each with duration d_n_/4 andfollowed by a break dr/12. Step III was obtained by applying the same transformation rule to each key press in step II thus obtaining the complete sequence [[K-2, K-1, K], [K-1, K, K+1], [K, K+1, K+2]].*

The typical trial was composed of three steps (I, II and III), and these steps unfolded according to one of three rules: Fractal, Iterative and Repetition (Figure 1). In steps I and II, participants executed the motor sequences as displayed visually on screen. The relation between steps I and II were determined by the underlying rule. Then, in step III, they were asked to generate a third motor sequence without visual guidance, corresponding to the correct continuation of step II using that particular rule.

The sequences comprising the first two steps were depicted visually on a computer screen by projecting colored circles onto the keys of a virtual keyboard. Different colors denoted the different fingers that participants had to use to press the keys (red: thumb, green: index and blue: middle finger). Thus, participants were asked to press the correct keys with the correct fingers, as demonstrated on the screen, synchronized both with the visual cues (coloured circles) and with the metronome beats. In these two first steps, participants were guided in the application of the rule. Then in step III, participants were asked to generate the correct continuation of step II, hence, to apply the rule one step further without the visual cues and following the temporal structure (Figure 1A).

The application of different rules always generated a final motor sequence of the kind [[K - 2*s*, K - *s*, K], [K - *s*, K, K + *s*], [K, K + s, K + 2s]]. To increase stimulus variability, we introduced the changeable parameters *s* and *k_0_*. The parameter *s* could be a value within the set {-2, -1, 1, 2}. If *s* was positive (1 or 2) the sequences were *ascending*, meaning that they unfolded from left to right on the keyboard (e.g. [K-1, K, K+1]). If *s* was negative (-1 or -2) the sequence was *descending*, meaning that sequences unfolded from right to left on the keyboard (e.g. [K+1, K, K-1]). When *s* =1 or -1, the sequence (within each cluster) was formed by *adjacent* keys ([K+1, K, K-1]), and when s = 2 and -2 the sequence was formed by *non-adjacent* keys, meaning that there was a space of one key between a pair of elements within the cluster (e.g. [K-2, K, K-2]). The initial key *k_0_* could be one of the middle four keys of the keyboard {7, 8, 9, 10}. Overall, these variations produced 16 different sequences, which were perfectly balanced across conditions. Crucially, these parameters had to be recognized during steps I and II, in order to correctly generate the sequence in step III.

### Pretest

All participants took part in a behavioral session up to one week before the fMRI experiment. The goal of this session was to instruct participants explicitly about the task rules, to assess their understanding of those rules and to train them in the execution of the motor sequences. Note that none of the participants had experience in playing music, particularly not in playing on a piano. The session lasted approximately 2 hours. Participants started by performing a beat perception task (part of the battery used in Müllensiefen et al., 2014) , to evaluate whether they were able to understand the synchrony between two temporal events (a metronome and a music piece). Then, they were shown a slideshow explaining the task rules, and video examples with the motor sequences they had to perform.

After the instructions, participants performed a supervised session comprising 10 trials following the Fractal rule. A researcher was in the room supervising this session, incentivizing the participants to follow the temporal structure and to use the visual landmarks to find the correct keys. Here, they were again asked to execute the sequences as depicted on the screen, but not asked to mentally generate the sequences by themselves. Steps I, II and III were all presented visually on screen (unlike in the task used in the MR scanner). The goal of this phase was to train the participants with correct exemplars. We repeated this procedure for the Iterative rule.

If participants were able to execute adequately the Fractal and Iterative sequences, they proceeded to a final session, similar to the one used in the fMRI. This session was composed of 20 trials including 8 trials following the Fractal rule, 8 trials following the Iterative rule, and 4 trials following the simple Repetition. In this last session, step III was not cued visually, and participants had to generate the sequence by themselves, without visual support, but following the metronome, as later in the MR scanner.

Accuracy was measured as the number of correct keys pressed at the correct time – within the interval [-.25s, 1s] locked to the onset of the appropriate beat. Each trial contained 9 expected key presses in step III.

Participants with accuracy >80% in the last session, for all rules, were invited to participate in the fMRI experiment. Out of 39 participants performing the pretest, only 21 fulfilled this criterion.

### fMRI Procedure

On the day of fMRI data acquisition, participants were again debriefed on the task rules, then positioned in the scanner and asked to perform a short test session of 6 trials. If they were able to perform adequately, we proceeded with the anatomical and functional data acquisition. Only one participant was excluded due to inability to replicate the experiment within the MR experimental apparatus. The whole procedure (briefing, scanning, and post testing) had a duration of approximately 2:30 hours.

Participants performed 4 task sessions, each with an approximate duration of 15min and composed of 20 trials - 8 ‘Fractal’, 8 ‘Iteration’ and 4 ‘Repetition’ trials. Trials following different rules were inter-mixed within each session and pseudo-randomized. The trial sequence was determined using Optseq2 (https://surfer.nmr.mgh.harvard.edu/optseq/) to maximize the efficiency of fMRI signal acquisition.

Trial structure is depicted in Figure 2. We were interested in two periods within each trial, namely the transition between step II and III – the *planning phase* – and in step III – the *execution phase.* The *planning phase* was important to capture the computations necessary to transform step II into step III, and the neural systems instantiating these transformations.

**Figure 2.**
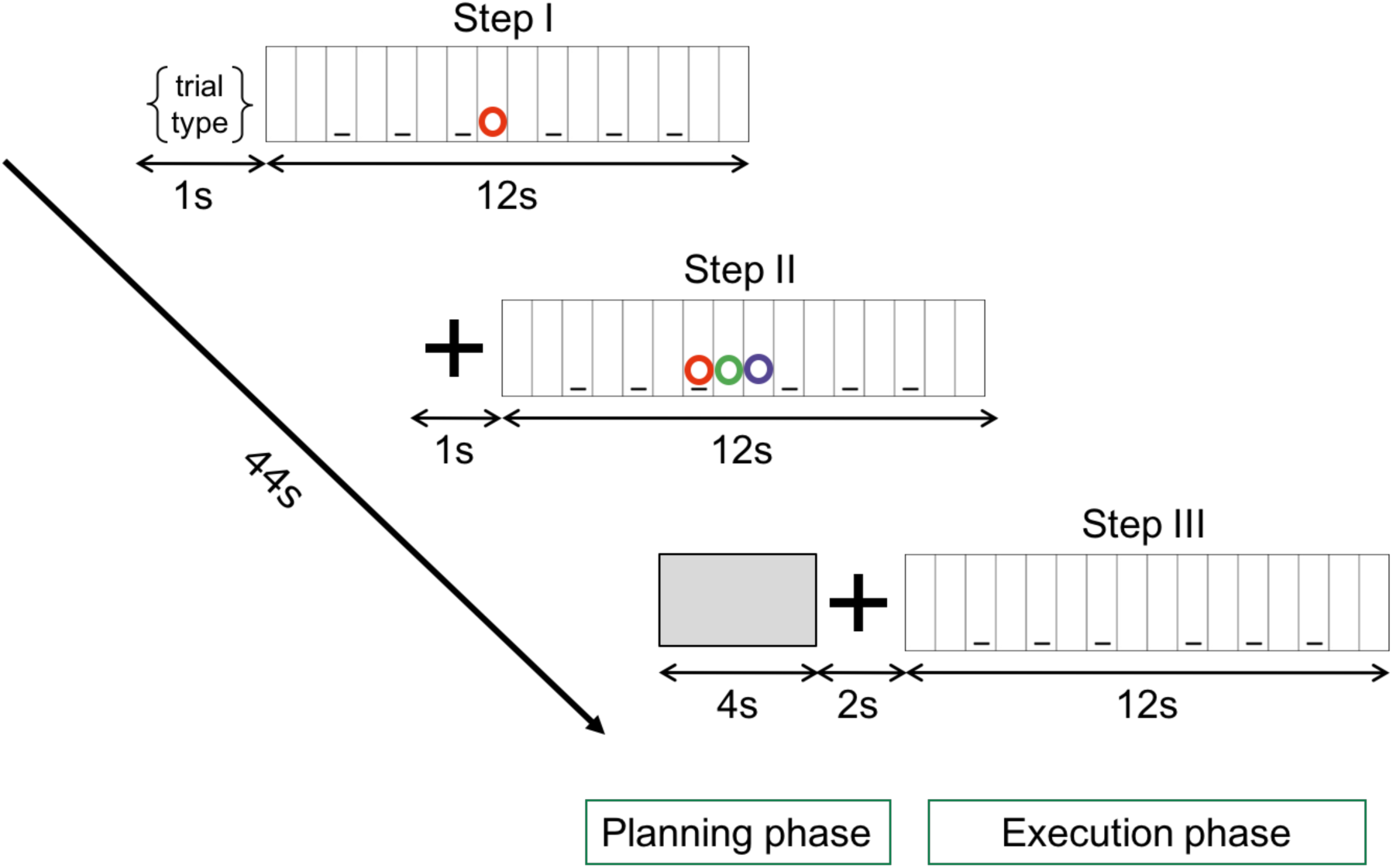
Trial structure. Inside the MR scanner (3-T Skyra, Siemens), participants performed 4 sessions of 20 trials each (8 Fractal, 8 Iteration and 4 Repetition trials). All trials had the same structure: First, a letter indicated the trial type (Fractal, Iteration or Repetition). Then, steps I and II of the sequence were shown on screen, which participants had to execute simultaneously on a keyboard (coloured circles indicated which finger to use). This was followed by a 6 seconds ‘planning phase ’ composed of a 4 seconds blank screen and a 2 seconds crosshair during which participants planned execution of step III. Finally, in the ‘execution phase ’, participants performed the correct continuation of the sequence without visual cues. Throughout all steps, a metronome sound at 60bpm guided participants ’ pace and the sequence’s temporal structure.

The experimental apparatus is depicted in Figure 3. Participants performed the task while lying in the MR scanner, using a silent 16-key MR-compatible piano (Figure 3A). We used a dual mirror system, so that participants were able to see both the virtual keyboard projected on the screen, and the physical keyboard on which they executed the motor sequences (Figure 3B). The position of the mirrors was adjusted individually for each participant. Both keyboards had visual markers on specific keys for visuo-spatial reference (Figure 3C). On the physical keyboard, these references could also be detected by touch. All participants used their right hand to perform the motor sequences.

At the end of the procedure, participants were given a questionnaire on their cognitive strategies used to generate step III across the different tasks (Supplementary Table S1).

**Figure 3.**
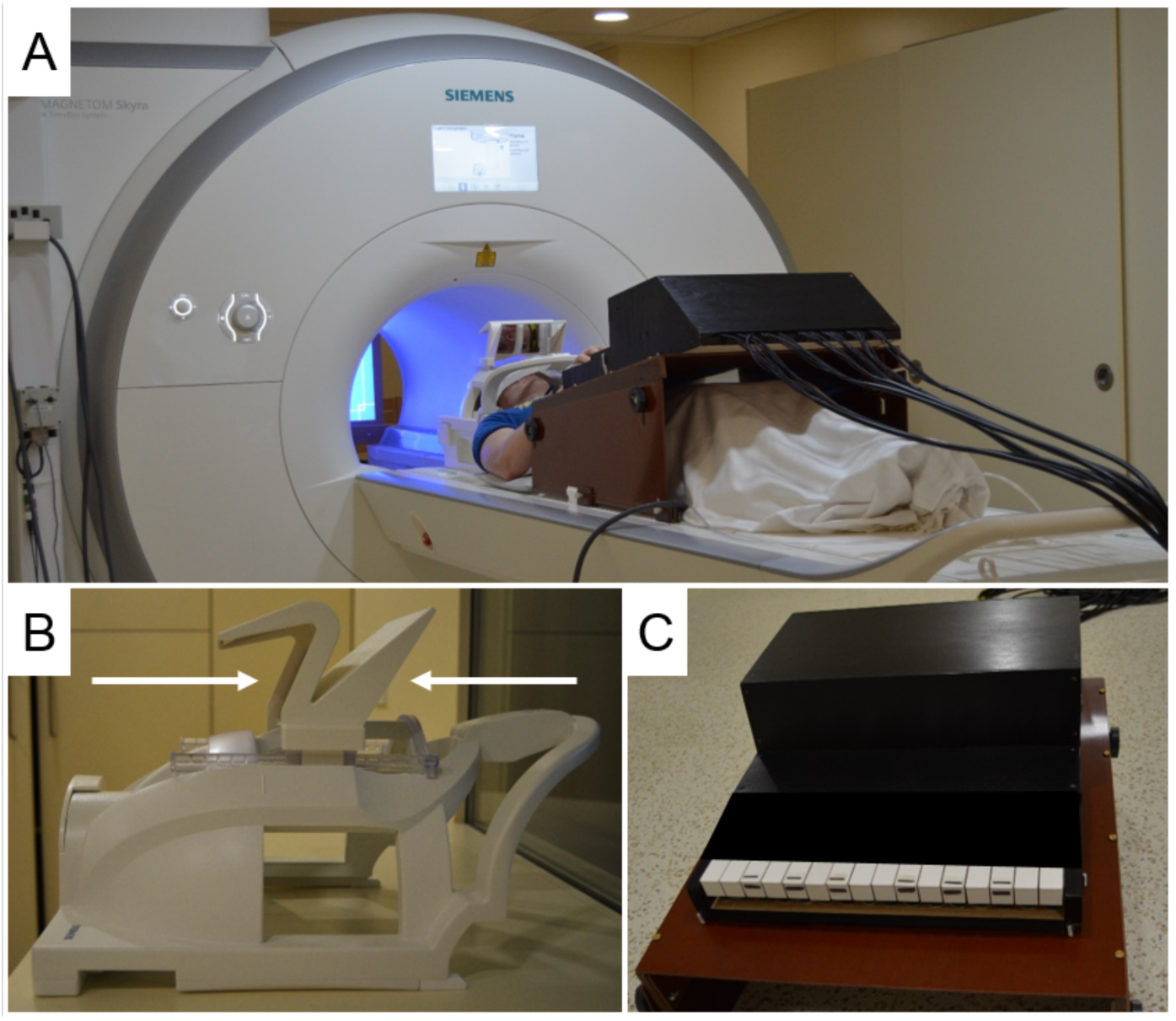
fMRI apparatus. **A.** The keyboard was placed on a custom-made wood stand. This stand provided a degree of inclination that increased the visibility of the keyboard. The metronome sound was delivered though MR compatible headphones. **B.** We used a double mirror system mounted on the head coil, which allowed participants to see both the virtual keyboard on screen (top mirror, left arrow), and the physical keyboard under their right hand (bottom mirror, right arrow). We adjusted the position of the mirrors for each participant to maximize visibility and comfort. **C.** The keyboard was an adapted MR compatible piano in which the black keys were covered. We added visual and tactile cues on specific keys that the participants could use for reference. Importantly, pressing the keys on the keyboard did not generate any sound, and therefore key-tone associations could not be used in our task, which was purely visuo-motor.

### Data Acquisition

The experiment was carried out in a 3.0-Tesla Siemens SKYRA whole body magnetic resonance scanner (Siemens AG, Erlangen, Germany) using a 32-radiofrequency-channel head coil. During the 4 sessions, functional magnetic resonance images were acquired using a T2*-weighted 2D echo planar imaging (EPI) sequence with TE = 30 ms and TR = 2000 ms. For each session, we acquired 450 volumes with a square FOV of 192 mm, with 31 interleaved slices of 3 mm thickness and 30% gap (3 × 3 × 3 mm^3^ voxel size) aligned to the AC-PC plane, and a flip angle of 90°. T1-weighted images were selected from the database of the institute for anatomical registration. Some participants had high-resolution T1-weighted images that were acquired using a 3D MP2RAGE sequence (TI_1_ = 700 ms, TI_2_ = 2500 ms, TE = 2.03 ms, TR = 5000 ms) with a matrix size of 240 × 256 × 176, with 1 mm isotropic voxel size, flip angle_1_ of 4°, flip angle_2_ of 8°, and GRAPPA acceleration factor of 3.

### Data Analysis

#### Task-based fMRI

fMRI data of 20 participants were analysed with statistical parametric mapping (SPM8; Welcome Trust Centre for Neuroimaging; http://www.fil.ion.ucl.ac.uk/spm/software/spm8/). Anatomical data from high-resolution T1-weighted images were obtained by masking uniform tissue-contrast image with the 2nd inversion image from the MP2RAGE sequence. Functional data were pre-processed by following standard spatial pre-processing procedures. They consisted of: slice time correction (by means of cubic spline interpolation method), spatial realignment, co-registration of functional and anatomical data. Then, we performed spatial normalisation into the MNI (Montreal Neurological Institute) stereotactic space that included resampling to 2 × 2 × 2 mm voxel size. Finally, data were spatially low-pass filtered using a 3D Gaussian kernel with full-width at half-maximum (FWHM) of 8 mm and temporally high-pass filtered with a cut-off of 1/128 Hz to eliminate low-frequency drifts.

Participants performed 4 sessions. For each of the 4 sessions (sess1-4), statistical parametric maps for the whole brain data were generated in the context of the general linear model (GLM) separately for the “planning” and the “execution” phase. The evoked hemodynamic response to the onset of one or the other phase was modelled for the fractal rule (F), iteration rule (I) and simple repetition (R) conditions as boxcars convolved with a hemodynamic response function (HRF). To this design, we added estimated motion realignment parameters as covariates of no interest to regress out residual motion artefacts and increase statistical sensitivity. For random effects group analyses, a within-subject flexible factorial ANOVA was performed on whole brain data with binary grey matter masks thresholded at intensity value of 0.25. A main effect of TASK was detected in both *planning* and *execution* phases, while a main effect of SESSION was present only in the *execution phase*. No interaction was detected. We created statistical parametric maps with t contrasts (SPM{T}). In the *planning phase,* t contrasts were calculated between each task with the following contrasts: F > I, F > R, I > F, I > R, R > I, R > F. We controlled family-wise error rate (FWER) of clusters below 0.05 with a cluster-forming height-threshold of 0.001. Anatomical labels are based on Harvard-Oxford cortical and sub-cortical structural atlas implemented in FSL (http://neuro.debian.net/pkgs/fsl-harvard-oxford-atlases.html) and for white matter fibres we used the JHU DTI-based white-matter atlases (http://neuro.debian.net/pkgs/fsl-jhu-dti-whitematter-atlas.html).

#### PPI

In the *execution phase,* we found common activation between the contrast F > I and R > I in a broad fronto-striatal cluster with main peak in the right uncinate fascicle (common maxima: x = 28, y =10, z = -12, cluster extent = 1080 voxels for the F > I contrast and 745 voxels for the R > I contrast). As post-hoc hypothesis, we tested whether the maximum common to the contrasts F > I and R > I interacted with motor regions during the implementation of F and R sequences. We thus examined the change in the effective connectivity of the inferior frontal cluster between F and I tasks, as well as R and I tasks using PPI analysis (Friston et al. 1997). Similar results in the PPI were obtained by both contrast-specific analyses. In order to extract voxels of interest (VOIs), a 5 mm sphere was defined around the coordinates (x = 28, y = 10, z = -12) that constituted the peak maxima of both second level contrasts F > I and R > I. In each participants’ data, the centre of the VOI was set to these coordinates. The first eigenvariate of the functional MRI signal changes of the VOI was extracted and its mean response was multiplied by a task regressor with information about the experimental conditions (F > I or R > I). This interaction term between source signal and experimental treatment was the first regressor in the PPI analysis. Additionally, the mean deconvolved source signal of the VOI and the task regressor were included in the model as covariates of no interest.

## Results

In the present fMRI study, participants generated sequences of finger movements in 3 steps (I, II, III) following one of three rules (Figure 1): (1) a linear Iterative rule, (2) a hierarchical Fractal rule, or (3) simple Repetition. In the first 2 steps (I, II) participants executed sequences as demonstrated on the screen. Then, they were asked to generate step III without visual support (Figure 2). We focused our fMRI analysis on the transition between step II and III, the ‘planning phase’ reflecting the generative act, and on step III, the ‘execution phase’ reflecting the externalization.

Overall, we found that during action planning, the generation of new hierarchical levels (in the Fractal condition) was supported by a bilateral network of brain areas involved in motor planning and imagery (Hétu et al. 2013; Hardwick et al. 2013), including somato-motor and premotor cortices, cerebellum, lateral occipital cortex (LOC), and left pallidum. Furthermore, executing the final sequence when formed by Fractal or simple Repetition rules entailed an interaction between right fronto-striatal and left hand-motor areas (M1), while Iteration recruited left hand-motor area only. Crucially, regions of interest (ROI) analyses within left and right IFG lent no evidence for involvement of this region in the generation of new hierarchical levels.

### Generation Of New Hierarchical Levels Is Supported By General Networks Of Motor Planning

By measuring brain activity in the ‘planning phase’, we sought to identify neural networks underlying the cognitive processes that are relevant for the transition between step II and III, i.e., the generative act as such. In Repetition, this process consisted in holding the full sequence of step II in memory until its repeated execution in step III (Figure 1B). In the Iterative rule, this process required the serial addition of one key press to each cluster within a fixed hierarchical level (Figure 1C), without generation of new levels. The Fractal rule entailed the generation of new hierarchical levels (Figure 1D) by recursively substituting each key press K_n_ with a new sequence of three key presses [K_n+1_ – *s*, K_n+1_, K_n+1_ + *s*]. The parameters of the transformation rule to be applied in each trial’s planning phase, *s* and initial key (*k_0_*), could be inferred from the transition between steps I and II (see Methods for details). Participants considered it equally easy to extract these parameters in Fractal and Iteration (*p* = .2), while it was overall easier in the Repetition condition, as expected (*Wilcoxon signed-ranks*: *p*’s < .05 compared to both Fractal and Iteration).

Whole-brain results of the planning phase are depicted in Figure 4 and Table 1. We found a clear dissociation between the Fractal rule and both Repetition and Iteration. The generation of new hierarchical levels in motor sequences using the Fractal rule was supported by a bilateral network known to be involved in motor learning (Hardwick et al. 2013), motor planning (Elsinger, Harrington, and Rao 2006) and imagery of motor sequences (Hétu et al. 2013). This network included the sensorimotor and premotor cortices, cerebellum and lateral occipital cortex (all clusters *p* < .05 FWE corrected) and was present in both contrasts ‘Fractal > Iteration’ and ‘Fractal > Repetition’. Thus, it was more active for the ‘Fractal rule’ condition. Additionally, we found the left Pallidum active in the contrast ‘Fractal > Iteration’ but not in ‘Fractal > Repetition’.

**Figure 4.**
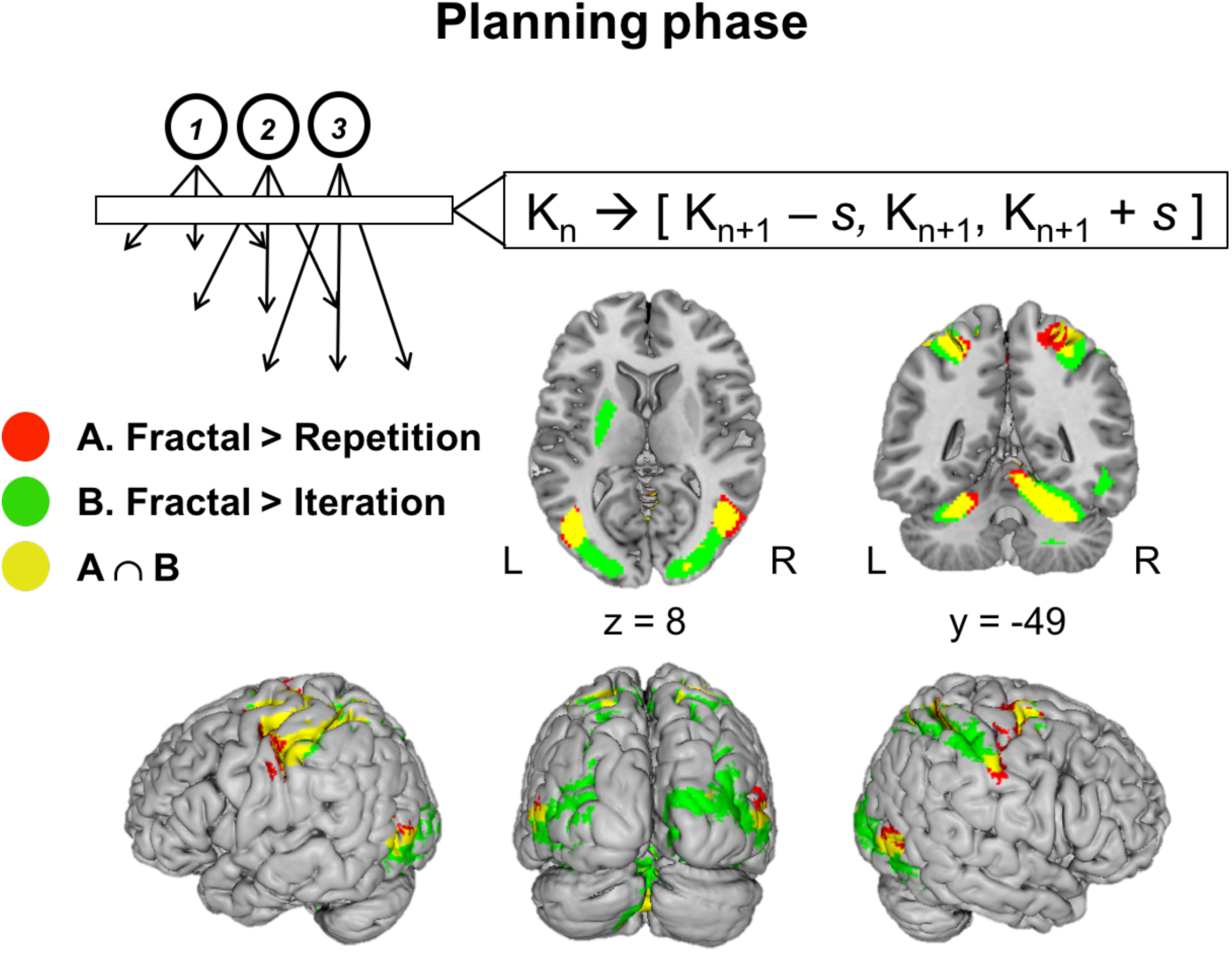
Brain activations during the planning phase (between steps II and III). Application of the Fractal rule yielded stronger activations compared to both Simple Repetition and Iteration in a bilateral network known to be involved in motor learning, planning and imagery, including sensorimotor and premotor cortices, cerebellum and lateral occipital cortex. The reverse contrasts (‘Iteration > Fractal’ and ‘Repetition > Fractal’) did not yield significant activations.

**Table 1.**
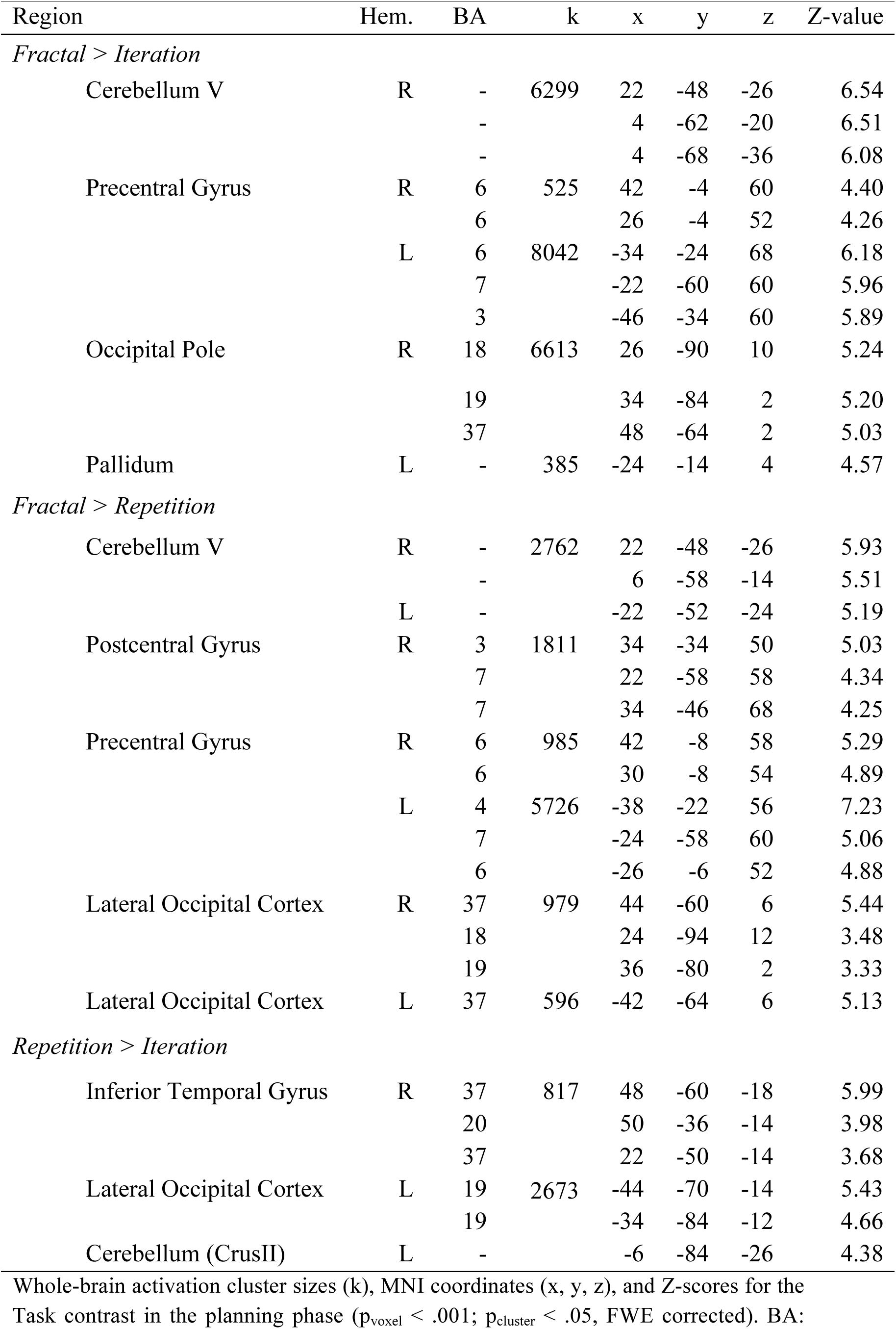
Task effect in the planning phase.

The dissociation between the Fractal rule and both Repetition and Iteration was nicely reflected in the subjective reports (see Supplementary table S1): in Fractal trials, participants (1) relied more on step II for the generation of step III than in both Iteration and Repetition (*p*’s < .05), (2) they imagined more where the hand should go in key space than in Iteration (z = -1.9, *p* = .059) and Repetition (z = -2. 0, *p* < .05), and (3) they more consciously prepared the sequence and thought about the rule in the Fractal condition than in Repetition (z = -2.1, *p* < .04). In line with these reports, no activations were found for the Iterative rule or Repetition, i.e., the contrasts ‘Iteration > Repetition’, ‘Iteration > Fractal’ and ‘Repetition > Fractal’ did not yield significantly active clusters. Only ‘Repetition > Iteration’ revealed activity in bilateral inferior lateral temporo-occipital cortex (Table 1).

To test whether there were specific activations for the Fractal rule within lateral PFC, particularly IFG, we performed four Small Volume Corrected (SVC) analyses within left Brodmann Area (BA) 44, left BA 45, right BA 44 and right BA 45. We found no significant differences between tasks (with uncorrected *p* < .01).

In sum, these findings are in line with the assumption that in the Fractal condition, participants engaged in specific computations to transform step II into the final step III sequence using an explicit motor-spatial rule. These computations were supported by general networks associated with motor planning and imagery, but did not specifically recruit IFG.

### Iterative Sequence Completion Is Solved by peripheral Sensorimotor resources

In the ‘execution phase’, we found clear similarities between Fractal rule and Repetition that both dissociated from Iteration (Figure 5 and Table 2). Note that this pattern of results cannot be explained by task difficulty because (1) the sequences to execute were motorically identical across all conditions, (2) key press accuracy did not differ between conditions (Fractal: mean ± *SD* = 87% ± 20%; Iteration: 89% ± 18%; Repetition: 87% ± 23%; Generalized score *χ2* = 1.8, *p* = .4), and (3) participants reported similar confidence in the correctness of their performance in Fractal and Iteration (*Wilcoxon signed ranks*: *p* > .4), and overall higher confidence in Repetition (*p*’s < .05).

**Figure 5.**
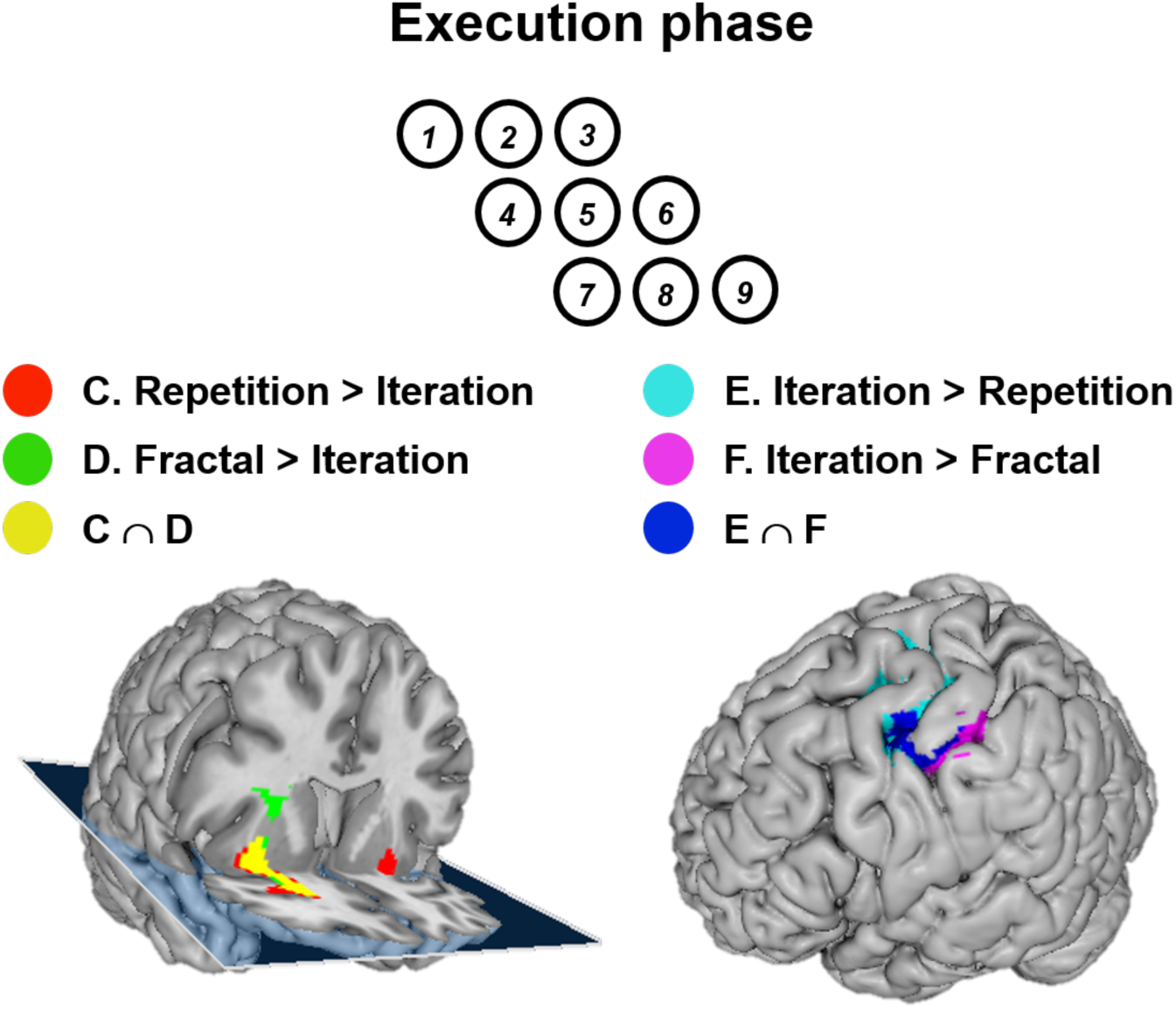
Brain activations during the execution phase (step III). Participants executed sequences of nine key presses that were identical at the motor output but generated according to different rules (Fractal, Iteration and Repetition). (1) In contrast with Iteration, both the Fractal rule and Repetition (C and D) activated a right fronto-striatal cluster (including the basal ganglia and orbitofrontal cortex, with peak in right uncinate fasciculus; x = 28, y = 10, z = -12). This cluster interacted with the hand area in left motor cortex (see details of PPI in text); (2) Conversely, execution of the same motor sequences using the Iterative rule (E and F) activated the left sensorimotor hand area only (x = -52, y = -18, z = 50).

**Table 2.** Task effect in the execution phase.

| Region | Hem. | BA | k | x | y | z | Z-value |
| --- | --- | --- | --- | --- | --- | --- | --- |
| *Iteration > Fractal* |  |  |  |  |  |  |  |
| Postcentral Gyrus | L | 4 | 1047 | -52 | -18 | 50 | 6.06 |
|  |  | 3 |  | -36 | -20 | 48 | 4.56 |
|  |  | 6 |  | -30 | -24 | 62 | 3.46 |
| *Fractal > Iteration* |  |  |  |  |  |  |  |
| Fronto-striatal<br>(peak in Uncinate Fasciculus) | R | 48 | 745 | 28 | 10 | -12 | 4.94 |
|  |  | 48 |  | 28 | 6 | 14 | 4.67 |
|  |  | 48 |  | 22 | 20 | 10 | 4.17 |
| *Repetition > Iteration* |  |  |  |  |  |  |  |
| Fronto-striatal<br>(peak in Uncinate Fasciculus) | R | 48 | 1080 | 28 | 10 | -12 | 5.36 |
|  |  | 48 |  | 32 | -2 | -18 | 4.58 |
|  |  | 47 |  | 26 | 34 | -8 | 4.53 |
| Amygdala | L | 34 | 2216 | -30 | -4 | -16 | 5.05 |
|  |  | 20 |  | -42 | -6 | -28 | 4.59 |
|  |  | 37 |  | -34 | -34 | -14 | 4.53 |

Whole-brain activation cluster sizes (k), MNI coordinates (x, y, z), and Z-scores for the Task contrast in the execution phase ( $p_{\text{voxel}} < .001$ ; $p_{\text{cluster}} < .05$ , FWE corrected). BA: Brodmann area, Hem.: hemisphere.

Sequence execution using the Iterative rule, compared to both Fractal rule and Repetition, yielded stronger activations in the sensorimotor hand area with a cluster peak in somatosensory cortex (x = -52, y = -18, z = 50 for both ‘Iteration > Fractal’ and ‘Iteration > Repetition’). Conversely, both contrasts ‘Fractal > Iteration’ and ‘Repetition > Iteration’ yielded significant activations in a large right fronto-striatal cluster which included putamen, amygdala, portions of orbitofrontal cortex and peaked in uncinate fasciculus (x = 28, y = 10, z = -12;*p* < .05 FWE corrected). A similar left-hemispheric cluster was found for ‘Repetition > Iteration’ only (peak in left amygdala: x = -30, y = -4, z = -16, extending into medial temporal lobe and putamen). No significant differences were found between Repetition and Fractal rule.

To test a post-hoc hypothesis of task-related functional connectivity of these fronto-striatal clusters with the sensorimotor hand area, we performed a whole-brain PPI analysis seeding in the maxima of the fronto-striatal cluster in the right hemisphere that was common to the two contrasts. Indeed, we found significant interactions with the hand area in left M1 (‘Repetition > Iteration’: x = -32, y = -26, z = 62, cluster extent = 1427 voxels, Z = 5.02, *p*_voxel_ < 001,*p*_cluster_ < 001; ‘Fractal > Iteration’: x = -36, y = -24, z = 48, cluster extent = 182 voxels, Z = 3.96, *p*_voxel_ < .001, *p*_cluster_ = .114).

Similar to the planning phase, we performed four Small Volume Corrected (SVC) analyses within left BA 44, left BA 45, right BA 44 and right BA 45. We found no significant differences between tasks (with uncorrected *p* <x .01).

In sum, during execution of identical motor sequences (that were, however, derived from different rules), we found cognitive and neural dissociations between Iteration and the other two rules. These effects were unrelated to the difficulty of execution. While the Iterative rule correlated more strongly with primary sensorimotor systems, both Repetition and Fractal rules involved additional motor control instantiated by fronto-striatal areas.

## Discussion

To our knowledge, the present study is the first to investigate the neural systems involved in the generation and overt production of motor hierarchies. In addition, we could separate the core generative act from its externalization components. To do so, we developed a novel paradigm that contrasted (1) sequences of finger movements formed according to a hierarchy-generating Fractal rule with (2) identical sequences formed according to rules that did not require generation of new hierarchical levels (Iteration and Repetition). Each motor sequence was composed of two initial steps (I and II) to establish the rules and their parameters that then had to be applied to freely generate the correct continuation of the sequence in step III. Repetition implied buffering of the given motor sequence [[K-2, K-1, K] [K-1, K, K+1] [K, K+1, K+2]] and Iteration required the completion of a pattern [[K-2, K-1, _ ] [K-1, K,_ ] [K, K+1,_ ]] using within-level transformations. Only the Fractal rule entailed the generation of new hierarchical levels through the recursive substitution of each finger movement K_n_ with a sequence of three finger movements [K_n+1_ - s, K_n+1_, K_n+1_ + s]. Accordingly, participants reported mostly for the Fractal condition that they relied on step II to consciously prepare the final sequence, and imagined the sequence prior to execution. Nevertheless, Fractal and Iterative conditions did not differ in correctness of their execution or in subjective reports of general difficulty. Therefore, our discussion focuses on the cognitive and functional interpretation of the different neural responses associated with these two particular rules.

Our first important finding was that the generation of new hierarchical levels using the Fractal (compared to Iterative) rule was supported by a general network of areas involved in motor learning, planning and imagery (Hétu et al. 2013; Hardwick et al. 2013; Elsinger, Harrington, and Rao 2006). This bilateral network included the sensorimotor and premotor cortices, cerebellum, lateral occipital cortex, and left pallidum. Interestingly, in this study, which focused on the motor domain, these generative processes did not seem to specifically recruit IFG, an area thought to play an important role in the processing of hierarchies across domains (Jeon 2014; Fitch and Martins 2014; Fadiga, Craighero, and D’Ausilio 2009).

Our second relevant finding was that execution of both Fractal sequences and simple Repetition (compared to Iterative sequences) involved right fronto-striatal regions that were coupled with the left hand-motor area. This activity may reflect the buffering of previously formed motor sequences (see below). In contrast, the execution of Iterative sequences involved the sensorimotor hand area only. Note that these differences occurred despite the fact that the motor sequences were identical in their execution.

We interpret these results within the framework of discrete sequence production models (Verwey 2001; Verwey, Shea, and Wright 2014). According to them, performance involves (1) sequence generation and motor loading during planning, followed by (2) fast execution of the motor buffer content by effector-specific motor processors. The generation of new hierarchical levels in the Fractal rule puts particular strain on stage (1), the planning of the final sequence. Unlike in Repetition and Iteration where the motor program is (partly) available already in step II, performers have to use their rule knowledge in the Fractal condition to construct the appropriate sequence continuation. Interestingly, they seem to do so without involving lateral PFC/IFG as proclaimed hierarchy processor but by means of general mechanisms of visuo-motor imagery and planning as shown by stronger activity in bilateral visuo-motor networks (Hardwick et al. 2013).

Once formed, these motor programs are buffered in fronto-striatal areas and sent to the hand area for execution as suggested by increased cortico-striatal connectivity in the PPI analysis. Our data speak for a similar buffering and transmission during Fractal and simple Repetition that does, however, not apply to Iteration. This latter finding together with generally less involvement of resources for planning suggest that motor sequences following Iterative rules might be completed via mechanisms supported by the sensorimotor hand area directly during execution. In other words, the motor buffer may simply hold the 6-key sequence of step II, thus imposing less resource demands on the motor buffer, while the ultimate 9-key sequence is completed on the fly during execution.

Overall, the differences between Fractal and Iteration reflect different mechanisms of motor sequence generation involving between-level vs. within-level hierarchical processing: while the generation of new hierarchical levels in the Fractal rule demands more planning resources, serial completion of motor sequences in the Iterative rule is resolved using mechanisms supported by the primary sensorimotor cortex. This suggests that iterative, serial generation of motor sequences might be the default strategy of motor production when hinging mainly on the primary sensorimotor cortex.

### Prior hypotheses: The Role of lateral PFC/IFG

Based on current views that IFG is involved in the processing of hierarchies across many domains (Fitch and Martins 2014; Fadiga, Craighero, and D’Ausilio 2009; Jeon 2014) and in line with models of a posterior to anterior gradient of lateral PFC for hierarchical organization of actions (Koechlin and Summerfield 2007; Badre 2008), we hypothesized lateral PFC, and particularly IFG, to be a likely candidate for the motor *generation* of new hierarchical levels in our Fractal rule condition. However, we did not find evidence for specific involvement of this area in the generation of new hierarchical levels. How can our results be reconciled with the previous literature?

On the one hand, the absence of lateral PFC activation in our task might indicate that this region is sensitive to hierarchies of action goals (or other non-motor contextual dependencies (Badre 2008)), rather than to transparent rules describing cross-level relations in motor hierarchies (i.e. inducible without prior instruction) as tested in our task. Alternatively, the resources necessary to *discriminate* hierarchical sequences may not completely overlap with those used for the *generation* of new hierarchical levels, in that discrimination recruits numerous additional cognitive mechanisms that are not relevant during generation but may well account for IFG effects. For example, building hierarchies from sequential input during *discrimination* also poses demands on resources required more generally for sequence encoding, buffering and template matching (Bornkessel-Schlesewsky et al. 2015; Fitch and Martins 2014), that may not be taxed to the same degree during the *generation* of hierarchical structures in the motor domain. Importantly, most discrimination studies found greater IFG involvement in material that drew strongly on these general resources, e.g., by using sequences that were violations (Molnar-szakacs, Iacoboni, and Koski 2005; Novick, Trueswell, and Thompson-Schill 2005; Bianco et al. 2016), had greater ambiguity (Vitello and Rodd 2015; Rodd et al. 2015), longer dependencies or posed higher demands on working memory than respective control sequences (Braver et al. 1997; Baddeley 2003). This makes it difficult to dissociate the contribution of specific hierarchical generativity and general cognitive control/sequence encoding processes to the observed IFG activations. Our design not only balanced the amount of required cognitive control across conditions (recall that final sequences were always correct, unambiguous and identical across conditions, although based on different rules); it also allowed us to study hierarchy processing stripped off general processes required for parsing temporally evolving sequences by specifically targeting hierarchy *generation* (in the planning phase). Consequently, the fact that we did not find evidence for lateral PFC involvement challenges the notion of multi-domain hierarchical generativity in IFG (Fadiga, Craighero, and D’Ausilio 2009; Fitch and Martins 2014) and rather argues for its more general function during encoding of structured sequences.

### Theoretical implications and future research

Humans are exceptional in their ability to generate hierarchical structures. However, the cognitive and neural mechanisms underlying this capacity remain speculative. To date, it is linguistics that provides the most concise/influential account for hierarchical generativity (Berwick and Chomsky 2015), by proposing that this capacity results from three components: (i) a core recursive ability which assembles syntactic structures, (ii) peripheral systems necessary for their encoding and externalization (conceptual-intentional and sensorimotor)^1^ and (iii) the interface between (i) and (ii). Within this framework, the core recursive ability (i) is hierarchical and independent of serial order. Thus, the apparent serial order of language would be the result of processes of externalization, given that the motor output system is obligatorily serial.

This proposed architecture poses a general empirical challenge, not only in language but also in other domains. As reviewed above, previous research investigating the processing of hierarchical structures focused on the *discrimination* of well-formed structures vs. violations (e.g., AABB vs. AABA), and/or structures with long-distance vs. local dependencies (e.g., AABB vs. ABAB). Crucially, these designs entail to contrast fMRI activity between stimuli that are different in both their deep structure (internal, hierarchical) and surface structure (external, serial). As a consequence, we cannot separate the resources that are active due to the hierarchical generative processes from those due to differences in the externalization components (e.g. working memory, top-down predictions in sequence parsing).

Our design acknowledges these difficulties and circumvented them by explicitly separating the generative act (planning phase) from the externalization (execution phase). Additionally, since the structures to be generated and executed were identical across conditions regarding their surface structure, we could keep the externalization confounds constant. Therefore, we provide new support for the classical generative architecture (Figure 5) with two important findings. First, the Fractal rule, which heavily hinges on the generation of new hierarchical levels (or branches), increased demands on the motor planning network, and not IFG/PFC. Second, the Iterative rule, which is traditionally serial, increased demands on the externalization components, namely the sensorimotor cortex. This implies that in the domain that we studied here, motor specific (rather than multi-domain) functional networks qualify as neural bases for different components of hierarchy generation, and that this ability in other domains can (and should) be scrutinised by taking its different components into account.

One last important theoretical issue: Even though our design isolates the “generation of new hierarchical levels”, this pertains to the *application* of well-learned hierarchical rules, independently on how these were acquired (Lungu et al. 2014). In this expertise stage, “rule application” might reflect either the execution of combinatorial computations specifically involved in the generation of new hierarchical levels, or simply the retrieval of previously formed hierarchical representations (Figure 5), which remain stored as “schemas” within the motor network (Wiestler and Diedrichsen 2013). Despite one linguistic study attempting to separate the *acquisition* and *application* phases of hierarchical rule processing (Opitz and Friederici 2003), more investigation is needed to dissociate the neural bases of hierarchical generativity “on the fly” or “schema” retrieval (Figure 5). Although our study was not designed to solve this issue, a necessary future step will be to test unexperienced participants to isolate the initial generative processes underlying structure building, without potential previously formed schemas present in long-term memory.

**Figure 5.**
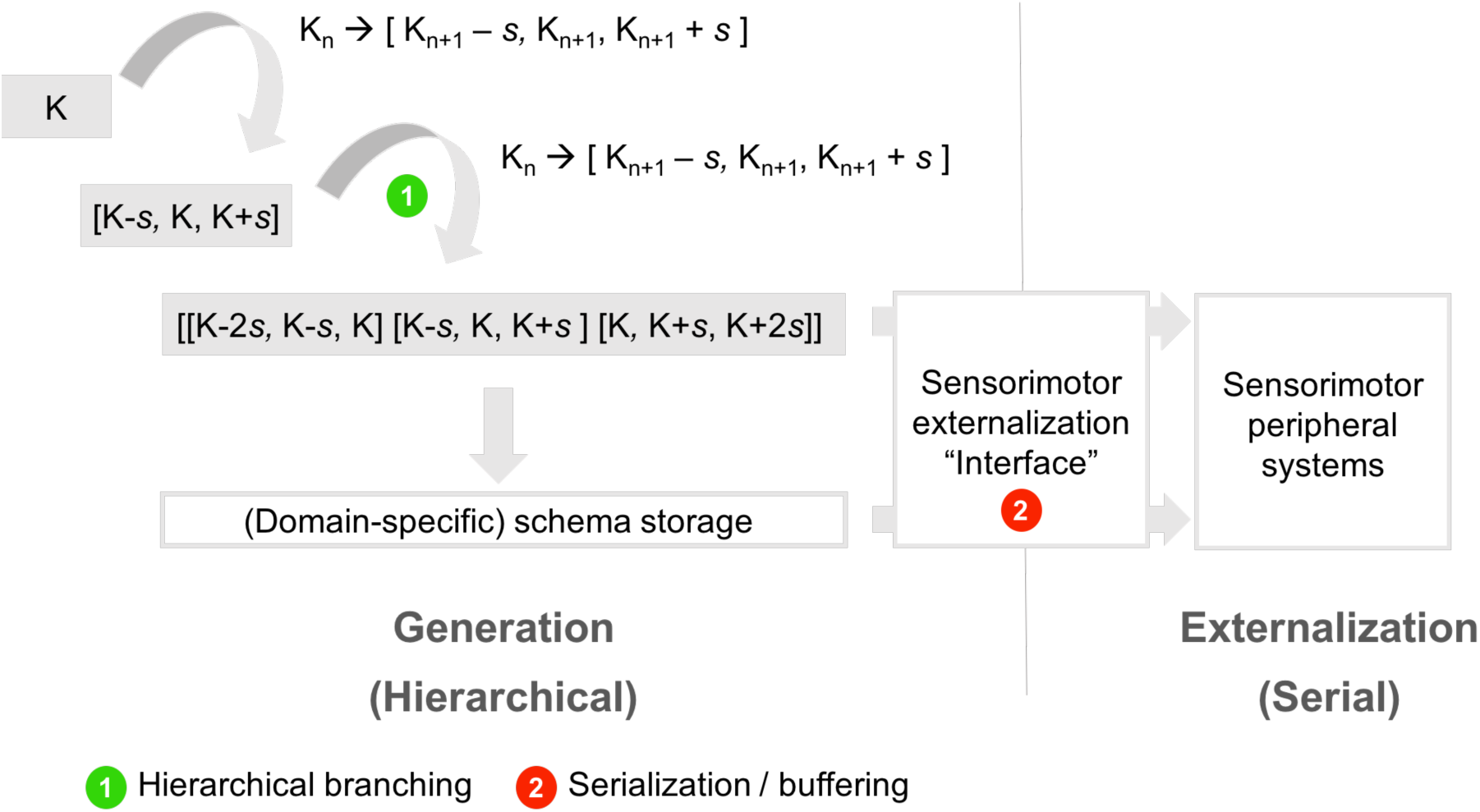
With our design, we explicitly separated the processes underlying the generation of hierarchical levels (left) from those used to externalize and execute motor programs (right). While the generation of new hierarchical levels in the Fractal rule involves hierarchical branching (left) which is not bounded to serial code, iterative completion of motor sequences is resolved using peripheral sensorimotor mechanisms (including computations within the primary sensorimotor cortices), which are serial (right). It should be mentioned that activations referring to the generation of new hierarchical levels can potentially involve either *de novo* combinatorial operations (upper cascade), or the retrieval of previously formed hierarchical representations (lower transparent box). The products of hierarchy-generating rules (e.g. [[K-2*s*, K-*s*, K] [K-*s*, K, K+*s*] [K, K+*s*, K+2*s*]]) might become schematized and stored in domain-specific networks from which they are retrieved during sequence generation. The schema would retain the clustered hierarchical structure and a set of free parameters (in this study the initial key K, and contour variable *s*). Importantly, even if the latter were the underlying mechanism, participants would have to extract and apply the parameters from the second step of each trial. Thus, irrespective of whether processing is based on combinatorial operations or retrieval of schemas, it would entail flexible generation of hierarchical motor sequences.

### Conclusion

In this study, we isolate the processes involved in generating motor hierarchies while separating them from other motor externalization components. Our results challenge the notion that a putative multi-domain hierarchical processor in the lateral PFC is necessary for the generation of hierarchical levels in motor sequence production. Instead, the generation of motor hierarchical structures via the application of recursive “fractal” rules was supported by a neural system used for motor learning and motor planning. While these systems might be important to *parse* hierarchical sequences in a multi-domain fashion, due to encoding and externalization processes, they do not seem to necessarily be so for the *generation* of new hierarchical levels.

## Acknowledgments

The authors are grateful to Sven Gutekunst and Jöran Lepsien for technical support.

## Author contributions

Mauricio J.D. Martins Roberta Bianco contributed project conception, experimental design and setup, data acquisition and analysis, data interpretation, writing the manuscript;

Daniela Sammler and Arno Villringer contributed supervision of the project, project conception, data interpretation, writing the manuscript.

## Data availability

The data that support the findings of this study are available from the corresponding author upon reasonable request. Authors can confirm that all relevant data are included in the paper and/or its supplementary information files.

## Footnotes

1 In the generative terminology, the primary sensorimotor cortex would be included in the “peripheral” encoding/externalization system, which is distinct from the anatomical notion of “peripheral” nervous system.

